# Translation elongation factor 1A2 (eEF1A2) is encoded by one of four closely related eef1a genes and is dispensable for survival in zebrafish

**DOI:** 10.1101/716753

**Authors:** Nwamaka J. Idigo, Dinesh C. Soares, Catherine M. Abbott

**Author notes:** Corresponding author: Prof Catherine M Abbott, Centre for Genomic & Experimental Medicine, MRC Institute of Genetics and Molecular Medicine, University of Edinburgh, Western General Hospital, Crewe Road, Edinburgh EH4 2XU, United Kingdom. ACS International Ltd., Begbroke House, Wallbrook Court, North Hinksey Lane, Oxford OX2 0QS, United Kingdom.

## Abstract

Zebrafish are valuable model organisms for the study of human single-gene disorders: they are genetically manipulable, their development is well understood, and mutant lines with measurable, disease-appropriate phenotypic abnormalities can be used for high throughput drug screening approaches. However, gene duplication events in zebrafish can result in redundancy of gene function, masking loss of function phenotypes and thus confounding this approach to disease modelling. Furthermore, recent studies have yielded contrasting results depending on whether specific genes are targeted using genome editing to make mutant lines, or whether morpholinos are used (morphants). *De novo* missense mutations in the human gene *EEF1A*2, encoding a tissue-specific translation elongation factor, cause severe neurodevelopmental disorders; there is a real need for a model system in which to study these disorders and we wanted to explore the possibility of a zebrafish model. We identified four *eef1a* genes and examined their developmental and tissue-specific expression patterns: *eef1a1l1* is first to be expressed whilst *eef1a2* is only detected later during development. We then determined the effects of introducing null mutations into eEF1A2 in zebrafish using CRISPR/Cas9 gene editing, in order to compare the results with previously described morphants, and with the severe neurodegenerative lethal phenotype of eEF1A2-null mice. In contrast with both earlier analysis in zebrafish using morpholinos and with the mouse eEF1A2-null mice, disruption of the *eef1a2* gene in zebrafish is compatible with normal lifespan. The resulting lines, however, may provide a valuable platform for studying the effects of expression of mutant human eEF1A2 mRNA.

## Introduction

Zebrafish represent a valuable model system for a range of human single gene disorders because they are genetically manipulable and their development is well understood. Importantly, if mutant zebrafish larvae have detectable phenotypes they can be used for high throughput small molecule library screening for the discovery of new therapeutically active molecules. However, redundancy of gene function as a result of gene duplication can confound this approach, and recent studies have yielded contrasting results that depend on whether specific genes are targeted using genome editing approaches like CRISPR/Cas9, or whether morpholinos are used (Law and Sargent, 2014; Kok *et al.*, 2015; Rossi *et al.*, 2015). One recently discovered human single gene disorder is *EEF1A2* related epilepsy, for which model systems are badly needed. In this study we sought to catalogue zebrafish *eef1a* genes, analyse their expression, and determine the effects of ablating expression of translation elongation factor eEF1A2 in zebrafish.

Translation elongation factor eEF1A, in its active GTP-bound form, is responsible for the delivery of aminoacylated-tRNAs to the acceptor site of the ribosome during the elongation step of protein synthesis. The elongation factor eEF1A is a member of the G protein family and is typically encoded by more than one gene, often located on distinct chromosomes in different eukaryotic species. Two sequence-redundant eEF1A genes TEF1 and TEF2 are present in the yeast *Saccharomyces cerevisiae* (Nagata et al., 1984; Nagashima, Nagata and Kaziro, 1986). In *Drosophila melanogaster*, two genes, F1 and F2, have been described (Hovemann et al., 1988) while four and five eEF1A genes has been reported in *Xenopus laevis* and *Solea senegalensis* respectively (Djé et al., 1990; Infante *et al.*, 2008; Newbery et al., 2011). In mammals, although numerous pseudogenes exist, only two active genes, *EEF1A1* and *EEF1A2*, encoding distinct but highly similar proteins (eEF1A1 and eEF1A2) have been reported (Ann *et al.*, 1992; Knudsen *et al.*, 1993; Chambers, Peters and Abbott, 1998; Kahns *et al.*, 1998; Svobodová *et al.*, 2015). These genes exhibit a developmental and tissue-specific pattern of expression: eEF1A1 is widely expressed during development but is then down-regulated in neurons, skeletal and cardiac muscle postnatally and replaced in these tissues with eEF1A2, which is concomitantly upregulated (Knudsen et al., 1993; Lee, Wolfraim and Wang, 1993; Chambers, Peters and Abbott, 1998; Svobodová et al., 2015).

In non-mammalian vertebrates the picture is less clear, but differential gene expression among the eEF1A genes has been noted for *Solea senegalensis* during larval development (Infante *et al.*, 2008). Expression of eEF1A genes in *Xenopus* is regulated post-transcriptionally. Newbery *et al.*, 2011 showed overlapping expression of eEF1A1 and eEF1A2 transcripts in the brain, heart and muscle tissues. However, at the protein level they observed a down-regulation of eEF1A1 in the brain and spinal cord and complete absence in *Xenopus* muscle. The eEF1A2 orthologue in *Xenopus* showed the same expression pattern as that of mammals, with expression restricted to the central nervous system and muscle tissues. While the importance of this isoform switching remains to be elucidated, it has been suggested that the isoforms may have additional distinct ‘moonlighting’ or non-canonical roles (reviewed in Ejiri, 2002; Mateyak and Kinzy, 2010) that are required for the different cell types (Abbott *et al.*, 2009).

There are several lines of evidence implicating translation elongation factor 1A2 (eEF1A2) in neurological disorders. A spontaneous deletion spanning 15.8 kilobases involving the promoter and first exon of *Eef1a2* is responsible for the wasted (*wst*) phenotype in mice (Chambers, Peters and Abbott, 1998; Newbery *et al.*, 2007). Mice homozygous for this mutation initially develop normally but then develop muscle wasting and neuronal degeneration from 21 days of age, the stage at which *Eef1a1* is down-regulated to undetectable levels in these tissues (Chambers, Peters and Abbott, 1998; Khalyfa *et al.*, 2001). The severity of the wasted phenotype progresses rapidly, leading to paralysis and death of the mouse by 28 days postnatal. On the other hand, heterozygous mice are healthy and do not show any muscular or neuronal abnormalities (Griffiths *et al.*, 2012).

More recently many heterozygous *de novo* missense mutations have been identified in individuals with neurodevelopmental disorders encompassing epilepsy, intellectual disability and autism (de Ligt *et al.*, 2012; Nakajima *et al.*, 2014; Veeramah *et al.*, 2014; Inui *et al.*, 2016; Lam *et al.*, 2016; Lopes *et al.*, 2016). Subsequently, Cao *et al.*, 2017 reported a homozygous missense *EEF1A2* mutation (P333L) in siblings that resulted in intractable seizures and death before the age of five from dilated cardiomyopathy. The severity of these disorders makes it important that model systems are developed for testing therapeutic strategies.

Zebrafish (*Danio rerio*) could provide a valuable model system for neurological disorders resulting from mutations in eEF1A2, but relatively little is known about eEF1A in zebrafish. In fact, it was first reported that only one eEF1A gene, which appeared to be developmentally regulated, was present in the zebrafish genome (Gao *et al.*, 1997). In addition, a gene identified as *eef1a* has been shown to be an essential gene required for early embryonic development in zebrafish (Amsterdam et al, 2004). More recently, Cao *et al*. 2017 reported that knockdown of *eef1a2* with morpholinos resulted in small head, cardiac failure and skeletal muscle weakness at 2 days post fertilisation (dpf). Together these results would suggest that mutation of any *eef1a* gene in zebrafish is lethal.

The complete sequence of the zebrafish genome is now available. Using this resource, we have identified and characterised the expression pattern of four *eef1a* genes; *eef1a1l1, eef1a1a, eef1a1b* and *eef1a2*, during development and in different adult tissues. We show *eef1a1l1* to be the embryonic form being the first to be expressed while *eef1a2* is the ‘adult’ form, detected later on during development. We went on to generate *eef1a2* null zebrafish using CRISPR-Cas9 gene editing and show in contrast with earlier analysis based on morpholinos, that disruption of the *eef1a2* gene is compatible with normal function in zebrafish.

## Materials and Methods

### Zebrafish husbandry and embryos and adult tissues collection

Zebrafish of the AB strain were used for all experiments. They were maintained in the MRC Human Genetics Unit (HGU) zebrafish facility at the University of Edinburgh according to standard procedures (Westerfield, 2000). Embryos were raised at 28.5°C in Petri dishes containing E3 embryo medium and staged according to Kimmel *et al* 1995. For RNA isolation, embryos and larvae were collected, rinsed with water, snap-frozen in dry ice and stored at −70°C until needed. Adult zebrafish were killed by immersing in excess Tricaine. Tissues were quickly dissected, placed in RNAlater® solution (Invitrogen) and stored at −70°C until RNA preparation. All procedures were performed in accordance with the Home Office regulations and the University of Edinburgh.

### Bioinformatics

Nucleotide and protein sequences of the eEF1A genes for zebrafish and other species were obtained from the Ensembl genome browser (https://www.ensembl.org/index.html). Multiple alignments of protein sequences were carried out using Clustal Omega (https://www.ebi.ac.uk/Tools/msa/clustalo/) and BoxShade v3.21 (https://embnet.vital-it.ch/software/BOX_form.html). A phylogenetic tree was constructed with the MEGA version 6 software (Tamura et al., 2013) and analysed using the maximum likelihood method based on the Poisson correction model (Zuckerkandl and Pauling, 1965). The tree with the highest log likelihood (- 2809.8575) is shown. The percentage of trees in which the associated taxa clustered together is shown next to the branches. Initial tree(s) for the heuristic search were obtained automatically by applying Neighbour-Join and BioNJ algorithms to a matrix of pairwise distances estimated using a JTT model, and then selecting the topology with superior log likelihood value. The tree is drawn to scale, with branch lengths measured in the number of substitutions per site. The reliability of each branch was assessed using 1,000 bootstrap replicates and reliable assignment values indicated. The analysis involved 24 amino acid sequences. All positions with less than 95% site coverage were eliminated. That is, fewer than 5% alignment gaps, missing data, and ambiguous amino acids were allowed at any position. There were a total of 462 positions in the final dataset.

### RNA extraction and RT-PCR analysis

Total RNA from adult fish tissues and approximately 50 embryos/larvae per developmental stage was isolated using TRIzol® (Invitrogen) and cleaned up with the RNeasy Mini Kit (Qiagen) with on-column DNase treatment using RNase-free DNase (Qiagen) according to the manufacturer’s instructions. RNA concentration and integrity were analysed using the Agilent 2100 Bioanalyzer. Full-length cDNA was then synthesised with a mix of random and oligo (dT) using the AffinityScript Multiple Temperature cDNA Synthesis Kit (Agilent Genomics) according to the manufacturer’s protocol. RT-PCR was carried out using primers for the zebrafish eEF1A genes with the Phusion High-Fidelity PCR master mix. Zebrafish *actb2* was amplified as an internal control. Primer sequences are shown in Table 1. Primers for *actb2* were also used to assess cDNA for genomic DNA contamination, with an additional amplicon of 684 bp seen if genomic DNA was present. Products were run on a 2% agarose gel.

**Table 1.**
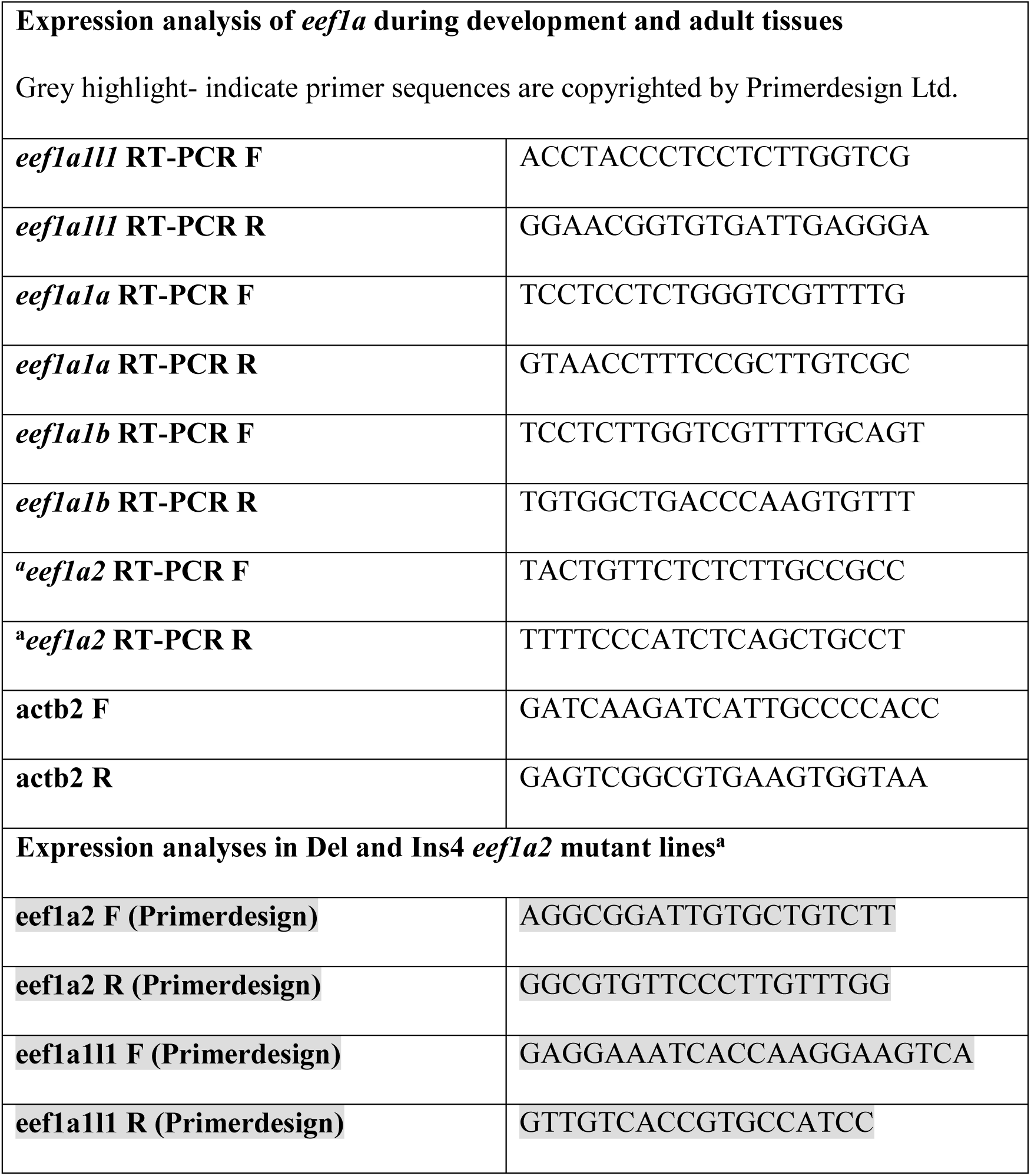
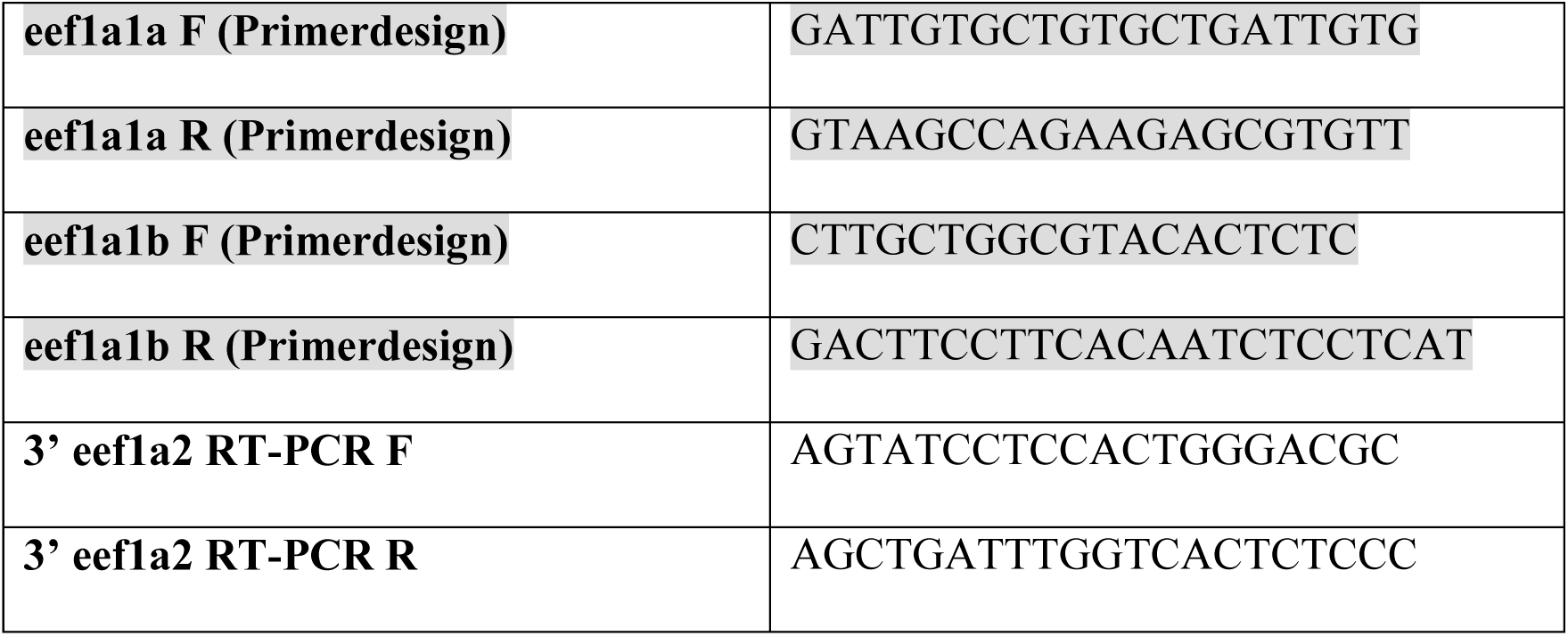
Sequences of primers.

### Quantitative real time PCR (qRT-PCR)

RNA and cDNA preparation from brain, muscle and liver tissues of adult fish was performed as described above. All qRT-PCR experiments were performed on diluted cDNA (1:5 in nuclease-free water) using the Brilliant II SYBR Green qPCR Master Mix (Agilent Technologies) and the 7900HT Real-Time PCR system (Applied Biosystems). 4 µl of each cDNA sample was added to 6 µl of qRT-PCR reaction mix following the manufacturer’s protocol. Reactions were performed under the following conditions: 95 °C for 10 minutes, 50 cycles of 95 °C for 30 seconds and 60 °C for 1 minute. For normalisation of gene expression, three reference genes were used: ATPsynth, NADH and 16S (selected using the geNorm kit from PrimerDesign Ltd UK). Prevalidated primers (PrimerDesign Ltd UK) were used for these experiments unless otherwise stated. Primer sequences shown in appendix table 1 are copyrighted by PrimerDesign Ltd UK. Sequences for the reference genes are however not disclosed by the company. To assess efficiencies, a standard curve was generated from seven 4-fold serial dilutions of pooled cDNA from whole adult fish (1:4, 1:16, 1:64, 1:256, 1:1024, 1:4096 and 1:16384) for each primer pair (Appendix table 2). Gene expression was quantified using the standard curve method. To compare the amount of each zebrafish *eef1a* transcript, the Pfaffl method (Pfaffl, 2001) was used to calculate the gene expression ratio of each target mRNA relative to the geometric average of the reference genes for each tissue. Results are based on the analysis of three biological replicates, each with triplicate technical replicates (a no-template control was included for each gene). Significance testing was performed using the Mann Whitney test or One-way ANOVA with Tukey multiple comparison tests where appropriate.

**Table 2:** Slope, intercept and correlation coefficient (R^2^) output from SDS software to estimate efficiency of primers used for qPCR analyses.

| **Primer** | **Slope** | **Y-Intercept** | **Correl. Coeff.**<br>**($R^2$)** |
| --- | --- | --- | --- |
| eef1a1l1 | -3.296 | 11.766 | 0.997 |
| eef1a1a | -3.223 | 19.571 | 0.997 |
| eef1a1b | -3.283 | 20.197 | 0.996 |
| eef1a2P | -3.294 | 21.767 | 0.994 |
| eef1a2S | -3.495 | 20.357 | 0.981 |
| 3' eef1a2 | -3.208 | 22.445 | 0.992 |
| ATPsynth | -3.237 | 14.832 | 0.999 |
| NADH | -3.227 | 16.500 | 0.996 |
| 16S | -3.018 | 10.194 | 0.991 |

### Expression vector construction and HEK293T cell line transfection

Total RNA and full-length cDNA were prepared from whole adult fish as described above. The zebrafish *eef1a* cDNAs were cloned into the destination vector, pcDNA6.2C-EmGFP for expression in mammalian cell lines using Gateway Cloning technology (Invitrogen) following the manufacturer’s instructions. The zebrafish *eef1a* genes were expressed together with a GFP tag in order to be able to discriminate between the exogenously expressed eEF1As and endogenous eEF1A1 of HEK293T cells. Each of the constructs were transfected into HEK293T cells using the TurboFect Transfection reagent (Thermo Fisher Scientific) according to the manufacturer’s protocol, with an empty vector control included. 24 hours before transfection, 2.4×10^4^ cells were seeded per well of a 6-well cell culture plate containing 4ml of DMEM (Gibco) with 10% Fetal Bovine Serum (FBS) growth medium. For each transfection reaction, 4 μg of plasmid DNA was diluted in 400 µl of serum-free DMEM (Gibco). The transfection reagent was briefly vortexed, then 6 µl was added to the diluted DNA. The reaction was mixed gently and incubated for 15 minutes at room temperature. The transfection mix (400 µl) was added to each well and mixed by rocking the plate gently. Cells were incubated at 37 °C in a CO_2_ incubator for 24 hours, after which they were analysed for transgene expression.

### Protein analysis

Protein lysates from adult zebrafish tissues and transfected HEK293T cells were prepared using RIPA lysing buffer with EDTA-free protease inhibitor (Roche). Concentration of protein lysates was determined using either the Pierce BCA protein assay kit (Pierce) or the DC Protein Assay (Bio-Rad) following the manufacturers’ instructions. Western blotting was carried out using near-infrared detection method by LICOR as previously described in Davies *et al*, 2017. Protein detection was also performed using the chemiluminescence method. In this case, after quantification of total protein using Sypro Ruby Blot Stain (Invitrogen) was analysed, blots were blocked at room temperature for 1 hour in 5% dried skimmed milk in TBS-0.1% Tween 20 (TBST). Membranes were incubated with primary antibody in blocking buffer overnight at 4°C, washed three times in TBST for 5 minutes and then incubated with the appropriate horseradish peroxidase-conjugated secondary antibody diluted in blocking buffer for 1 hour at room temperature. Finally, membranes were washed three times for five minutes each in TBST and protein detected using Clarity western ECL substrate (Bio-Rad) according to the manufacturer’s instructions.

### CRISPR/Cas9 experiment

Single guide RNA (sgRNA) targeting the zebrafish *eef1a2* gene was designed using the online tool CHOPCHOP (http://chopchop.cbu.uib.no/) and the oligonucleotides TAGGATAAGTTGAAGGCTGAGA and AAACTCTCAGCCTTCAACTTAT purchased from Integrated DNA Technologies (IDT) with a 5’ phosphate modification to increase ligation efficiency. The sgRNA construct was made by inserting annealed pairs of oligonucleotides into Bsal (New England Biolabs) digested pDR274 (Addgene #42250, Hwang *et al*, 2013) backbone. The single guide RNA plasmid was used as a template to amplify gRNA sequences, which were then transcribed using the Ambion MAXIscript T7 kit (Thermo Fisher Scientific). Cas9 mRNA was synthesised by transcribing NotI-digested pCS2-nCas9n (Addgene #47929, Jao *et al*, 2013) using the SP6 mMESSAGE mMACHINE kit (Thermo Fisher Scientific) to generate capped mRNA. Purification of synthesised mRNA was performed using SigmaSpin sequencing reaction clean-up kit (Sigma Aldrich) according to the manufacturer’s instructions.

#### Microinjection

Injection mixture containing 300 ng/µl Cas9 mRNA and 92 ng/µ sgRNA was prepared and injected into the cell of one-cell-stage zebrafish embryos. At 2 days post fertilisation (dpf), genomic DNA was extracted from a pool of 5-10 microinjected healthy embryos and the target region was amplified using the primer set 5’ CACCTTTATTTTTGCGTGAACA and 5’ TCAAAAACATGATCACTGGGAC. In order to assess the mutagenic efficiency of the gRNA, PCR products were TOPO cloned using the TOPO-TA cloning kit (Invitrogen) and individual clones were sequenced. Founder fish (3 months) were screened by amplifying target region using genomic DNA extracted from tail fin clippings and analysing the amplicons on the Agilent 2100 Bioanalyser.

#### Establishing stable mutant lines

Putative founders were outcrossed with wild-type AB fish. To determine whether the Cas9-induced mutations were heritable, genomic DNA from 10 individual embryos were assessed for indels, while the others were raised to adulthood. Mutant alleles were identified in F1 fish by Sanger sequencing and fish with identical mutations were mated. Confirmation of the sequence of the alleles was achieved with the homozygous F2 fish using Phusion High-Fidelity DNA Polymerase (NEB) and Sanger sequencing.

### Histology

Adult zebrafish from both *eef1a2* mutant lines and age-matched wild-type controls were fixed in 10% neutral buffered formalin (Sigma-Aldrich). Spinal cord sections were cut at a thickness of 3 µM. Paraffin-embedded spinal cord sections were dewaxed with xylene and rehydrated through a decreasing series of ethanol. Antigen retrieval was carried out using Proteinase K for 10 minutes at room temperature and slides were then treated with 3% hydrogen peroxide to block endogenous peroxidase. Sections were blocked with goat serum (1:5 in PBS) for 10 minutes. They were incubated overnight with anti-GFAP rabbit antibody (Dako) diluted at 1:500 in PBS, washed twice with PBS for 5 minutes and then incubated with anti-rabbit biotinylated antibody (Dako) at a concentration of 1:500 in PBS for 30 minutes at room temperature. Sections were then treated with Strept ABC reagent (Vector Laboratories) for 30 minutes and with Diaminobenzidine (DAB; Abcam) for 10 minutes. Sections were counterstained in haematoxylin solution (Shandon), dehydrated and finally mounted in DPX (VWR).

### Data availability

The authors state that all data necessary for confirming the conclusions presented in the article are represented fully within the article. All fish lines and plasmids are available upon request.

## Results

### Characterisation of the zebrafish eef1a genes

#### Bioinformatics analysis of eef1a genes in the zebrafish

To determine the full complement of *eef1a* genes in zebrafish, we identified four *eef1a* genes using Ensembl, namely, *eef1a1l1, eef1a1a, eef1a1b* and *eef1a2* located on chromosome 19, 13, 1 and 23 respectively. Each contained eight exons, seven introns and an open reading frame encoding distinct proteins of 462 (Eef1a1l1, Eef1a1a and Eef1a1b) or 463 (Eef1a2) amino acids. They also had similar exon-intron organisation with the 5’UTRs extending into exon 2 and the 3’UTR starting in exon 8. Whilst coding exons were of a consistent size, some introns of *eef1a1a, eef1a1b* and *eef1a2* are greatly expanded compared to the compact introns of *eef1a1l1* (Fig.1a). The zebrafish *eef1a* genes shared high sequence similarity at the nucleotide level within the coding region and also at the amino acid level, with *eef1a1a* and *eef1a1b* being particularly closely related (Fig. 1b)

**Fig. 1.**
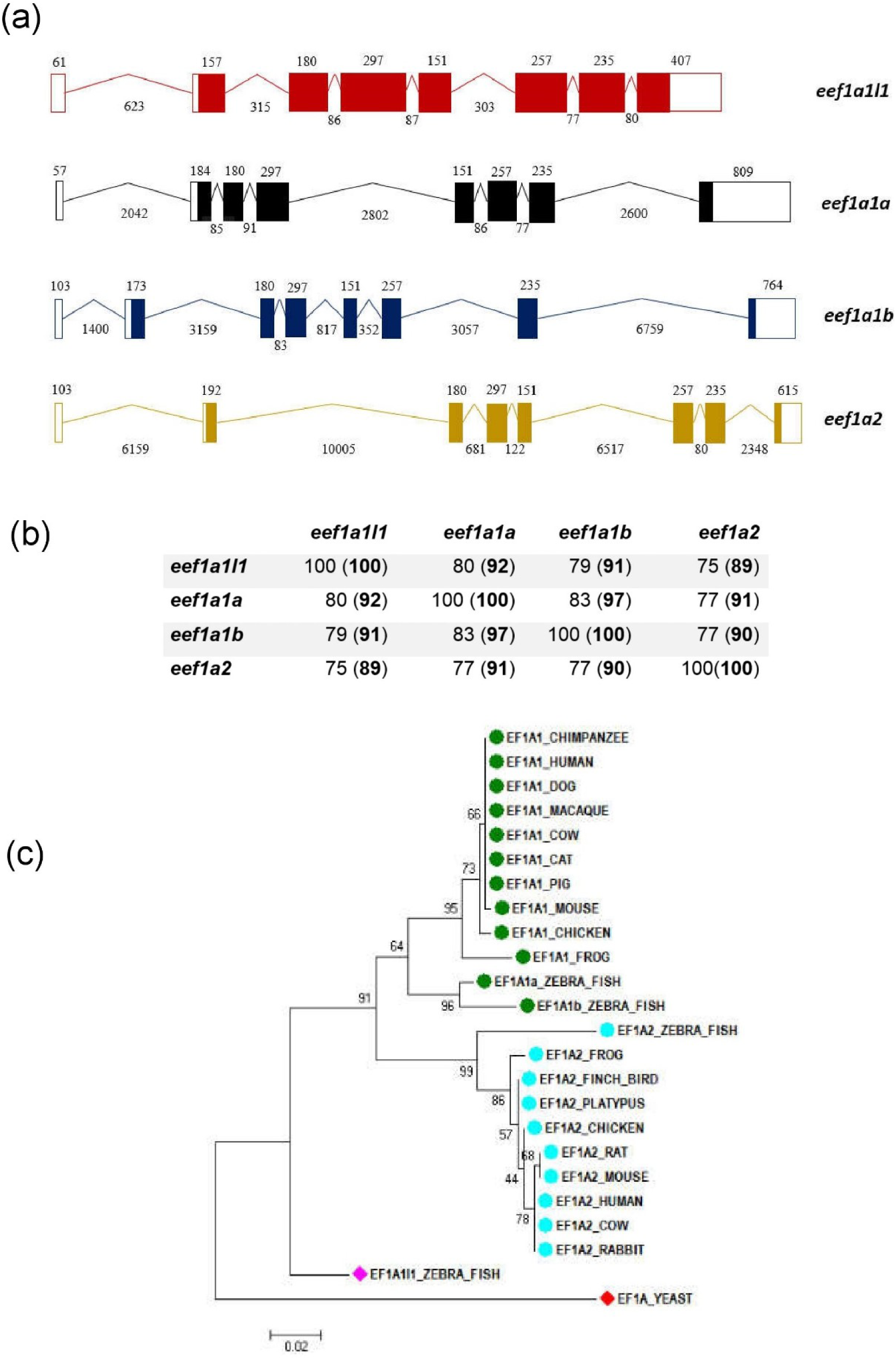
Four *eef1a* genes identified in the zebrafish genome. **a** Schematic representation of the exon-intron organisation of *eef1a1l1* (red), *eef1a1a* (black), *eef1a1b* (blue) and *eef1a2* (yellow) structures obtained from the Ensembl database. Length (in base pairs) of exons and introns, which are not drawn to scale, are indicated above and below respectively. **b** Percentage identity matrix for zebrafish eEF1As at the nucleotide and amino acid sequence level (in brackets) calculated using Clustal Omega. **c** Phylogenetic relationship among the zebrafish eEF1As and other vertebrate eEF1As using the Maximum Likelihood method.

The phylogenetic relationships of the four zebrafish Eef1a protein sequences and those from other vertebrate species were analysed using the maximum likelihood method (Fig. 1c). The phylogenetic tree obtained from this analysis showed both Eef1a1a and Eef1a1b to fall into the eEF1A1 clade while the zebrafish Eef1a2 segregated with eEF1A2 from the other vertebrates. In contrast, Eef1a1l1 did not cluster with any of the well-supported clades but appears to possess sequence features similar to both eEF1A1 and eEF1A2.

Alignment of the protein sequences of eEF1A protein sequences from zebrafish, mouse and human using Clustal omega (Fig. 2) also showed Eef1a1a and Eef1a1b to have higher sequence identity of ∼95% with mouse and human eEF1A1 than those obtained for Eef1a1l1 and Eef1a2. The zebrafish Eef1a2 showed higher sequence identity of ∼94% with mouse and human eEF1A2 than those obtained for Eef1a1l1, Eef1a1a and Eef1a1b. On the other hand, Eef1a1l1 had similar sequence identity to the eEF1A orthologues in mouse and human, with ∼92% for eEF1A1 and ∼90% for eEF1A2 in both species.

**Fig. 2.**
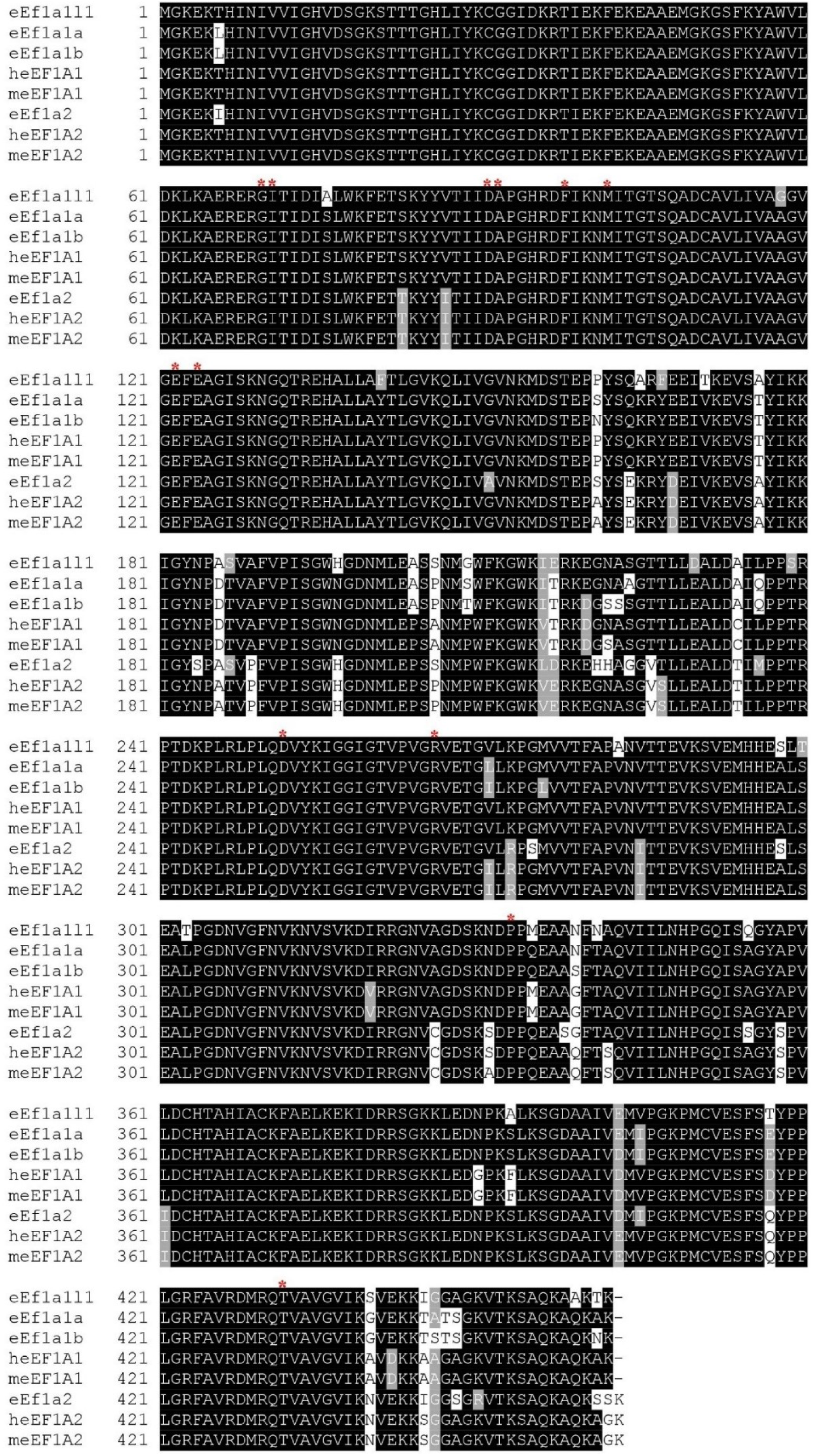
Multiple amino acid sequence alignment of eEF1A orthologues from zebrafish, human and mouse. The human eEF1A: heEF1A1 and heEF1A2, mouse eEF1A: meEF1A1 and meEF1A2. Identical and similar amino acid residues are indicated by black and grey backgrounds. Red asterisks (*) indicate some of the clinically important human eEF1A2 mutations, each of which involves residues that are completely conserved in the four zebrafish eEF1A isoforms (de Ligt *et al.*, 2012; Nakajima *et al.*, 2014; Veeramah *et al.*, 2014; Inui *et al.*, 2016; Lam *et al.*, 2016; Lopes *et al.*, 2016).

#### Expression of eef1a genes during development and adult tissue

The expression of each zebrafish *eef1a* gene was analysed at each of twelve different developmental stages; 1-cell, 2-cell, 4-cell, 8-cell, 16-cell, 256-cell, high, 50%-epiboly, 90%-epiboly, 24 hours post-fertilisation (hpf), 48 hpf and 72 hpf stages (Fig. 3a). Only *eef1a1l1* transcripts were detected at all the stages examined. Expression of *eef1a1a* and *eef1a1b* transcripts were detected at the 24 hpf, 48 hpf and 72 hpf developmental stages. The zebrafish *eef1a2* gene was the last to be expressed, being detected only at 48 and 72 hpf (Fig. 3b).

**Fig. 3.**
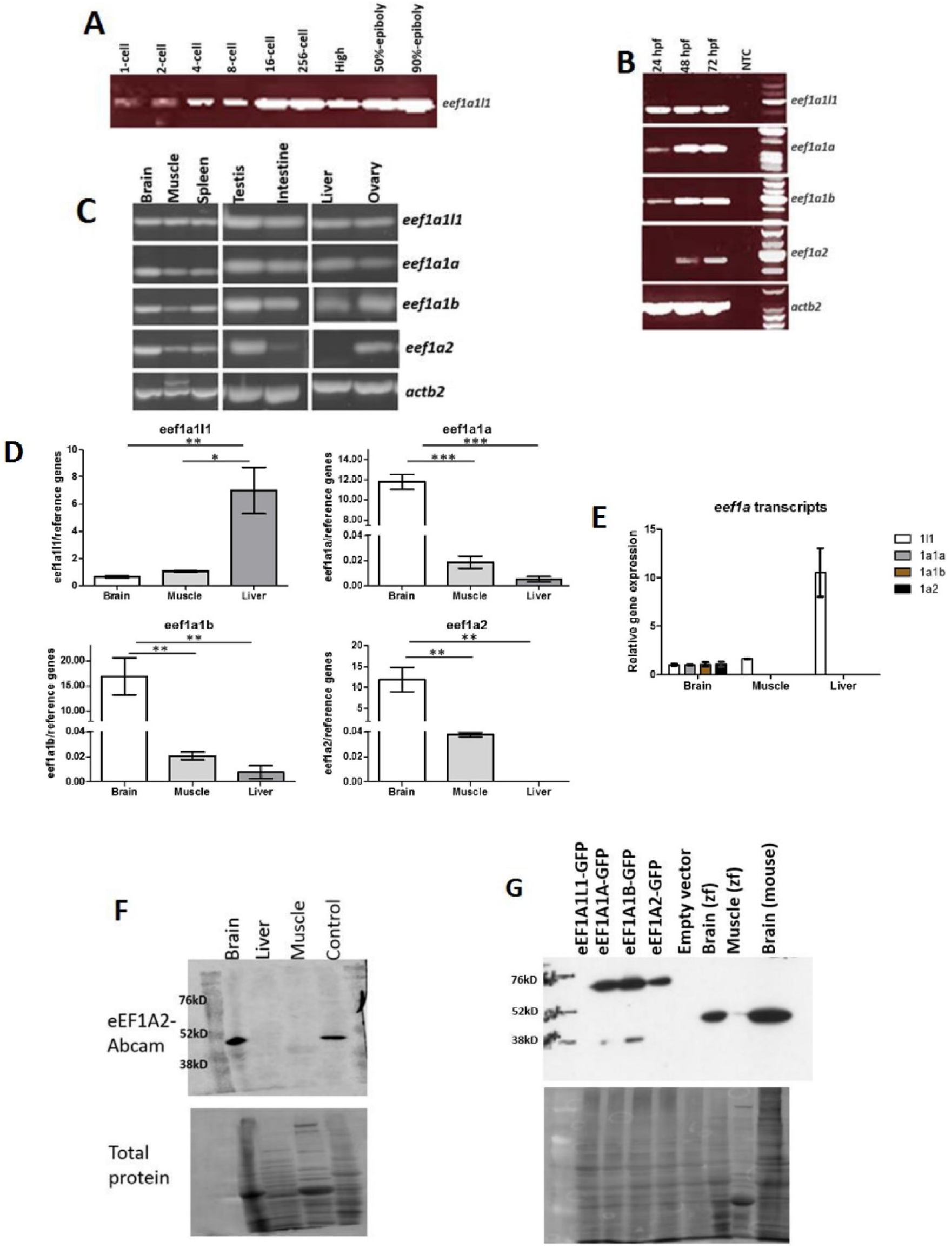
Expression analysis of the zebrafish *eef1a* genes. **a** Expression of *eef1a1l1* in different early embryonic stages detected by RT-PCR. The other eef1a genes were undetected at these stages (data not shown). **b** Expression of *eef1a* genes in 24 hpf, 48 hpf and 72 hpf developmental stages using RT-PCR. NTC – no template control. **c** Expression of *eef1a* genes in different tissues of adult zebrafish detected by RT-PCR. **d** Expression levels of *eef1a* genes in brain, muscle and liver tissues **e** and comparison of the relative levels of their transcripts in these tissues. Expression values were normalised to those of *ATPsynth, NADH* and *16S*. Results are means + S.E.M, n=3. *p < 0.05; **p < 0.01; ***p < 0.0001. For comparison, data were presented as the gene expression ratio of the target mRNA to the geometric mean of reference genes for each tissue. **f** Western blot showing Eef1a2 expression in brain, liver and muscle zebrafish tissues using eEF1A2-Abcam antibody (1:1000). Control is muscle tissue from mouse. **g** Validation of eEF1A2-Abcam antibody specificity using lysates isolated from individually transfected HEK293T cells with GFP-tagged Eef1a constructs. Zf-zebrafish.

We used RT-PCR to examine the expression of the zebrafish *eef1a* genes in adult tissues using total RNA extracted from brain, muscle, spleen, testis, intestine, liver and ovary (Fig. 3c). Three of the zebrafish *eef1a* genes, *eef1a1l1, eef1a1a* and *eef1a1b* were readily detected in all the tissues examined. Expression of zebrafish *eef1a2* was detected in brain, muscle, spleen, testis and ovary tissues but was only just detectable in the intestine. No *eef1a2* expression was seen in the liver. Using qPCR, we then analysed the level of expression for each *eef1a* gene in the brain, muscle and liver (Fig. 3d). The expression level of *eef1a1l1* was significantly higher in liver than in brain (p<0.01) and muscle (p < 0.05). Similar expression were seen for *eef1a1a, eef1a1b* and *eef1a2* with brain showing the highest level, followed by muscle and liver. The relative amount of the four *eef1a* transcripts in these tissues were calculated (Fig. 3e). In general, the most abundant transcript was that of *eef1a1l1* with approximately 7,980, 7,830 and 240-fold higher overall expression ratios than *eef1a1a, eef1a1b* and *eef1a2* respectively. While the relative amount of all the *eef1a* transcripts was similar in brain, *eef1a1l1* showed the highest value in muscle (1,040, 1,280 and 490-fold higher than *eef1a1a, eef1a1b* and *eef1a2* respectively) and liver (22,900 and 22,200-fold higher in *eef1a1a* and *eef1a1b* respectively) tissues. The relative amount of *eef1a2* transcripts in muscle was approximately two and three-fold higher than that of *eef1a1a* and *eef1a1b* respectively, both of which showed similar amounts in all three tissues examined.

#### Validation of commercially available eEF1A2 antibody

To investigate whether the *eef1a* mRNAs detected are translated into stable proteins we needed to identify a suitable antibody. Three commercially available antibodies against Eef1a2 were tested on protein lysates from adult zebrafish brain, liver and muscle tissues. The Genetex and Proteintech anti-eEF1A2 antibodies detected a band in liver (data not shown) in spite of the absence of Eef1a2 at the mRNA level (Fig. 3 c and d). However, an antibody from Abcam detected a band only in lysates from adult zebrafish brain and not in liver, consistent with our qRT-PCR data for Eef1a2 (Fig. 3f). This antibody was then used to test a range of other adult tissues: muscle, spinal cord, intestine, ovary and heart. Interestingly, expression was only detected in spinal cord and not muscle, in contrast to our RT-PCR results (data not shown).

Since the interpretation of results using this antibody could be complicated by the presence of other Eef1a paralogues, we carried out an antibody validation test. GFP-tagged constructs were made for each of the four zebrafish *eef1a* genes, transfected into HEK293T cells and lysates analysed with the eEF1A2-Abcam antibody. A band of the expected size was observed in lanes containing lysates isolated from cells transfected with GFP-tagged Eef1a1a, Eef1a1b and Eef1a2 (Fig. 3g), suggesting that this antibody cross-reacts with other Eef1a paralogues.

### CRISPR/Cas9 generated Eef1a2-null zebrafish survive to adulthood

We next sought to investigate the effect of loss of Eef1a2 in zebrafish using the CRISPR/Cas9 system. We wanted to establish whether zebrafish, like mice, undergo fatal neurodegeneration in response to the loss of eEF1A2 after developmental down-regulation of eEF1A1; consistent with the lethal phenotype arising from the use of morpholinos against eEF1A2 as shown by Cao *et al.*, 2017 or whether the presence of three paralogues in zebrafish gives rise to redundancy. A single gRNA targeting the zebrafish *eef1a2* gene was designed and microinjected, together with Cas9 mRNA, into one-cell embryos. The gRNA showed a mutagenic activity rate of ∼77% and a survival rate of 93% was observed within the CRISPR-injected embryos. At 2 months, adult (F0) injected zebrafish were genotyped using genomic DNA from tail fin clipping in order to identify potential founders. PCR amplicons containing the target region were analysed on the Agilent 2100 Bioanalyser which showed several distinct mutations at the target site were present in these fish (supplementary figure 1). In order to establish stable null mutant lines, F1 embryos were obtained from one of the mosaic putative founders outcrossed with wild-type fish. These embryos were then raised to adulthood and three mutant alleles, a 12 base pair deletion, a 4 base pair insertion and a 2 base pair deletion, were recovered (Fig. 4a). The 4 base pair insertion (hereafter referred to as Ins4) and 2 base pair deletion (hereafter referred to as Del2) were chosen for further analysis since they were each predicted to give rise to frameshifts resulting in premature stop codons (Fig. 4a). Heterozygous F1 zebrafish carrying the same mutations were then intercrossed and the embryos raised to adulthood. Interestingly, homozygous Ins4 and Del2 fish survived to adulthood with no obvious phenotypic differences from their wild type siblings, and were fertile (Fig. 4b).

**Fig. 4.**
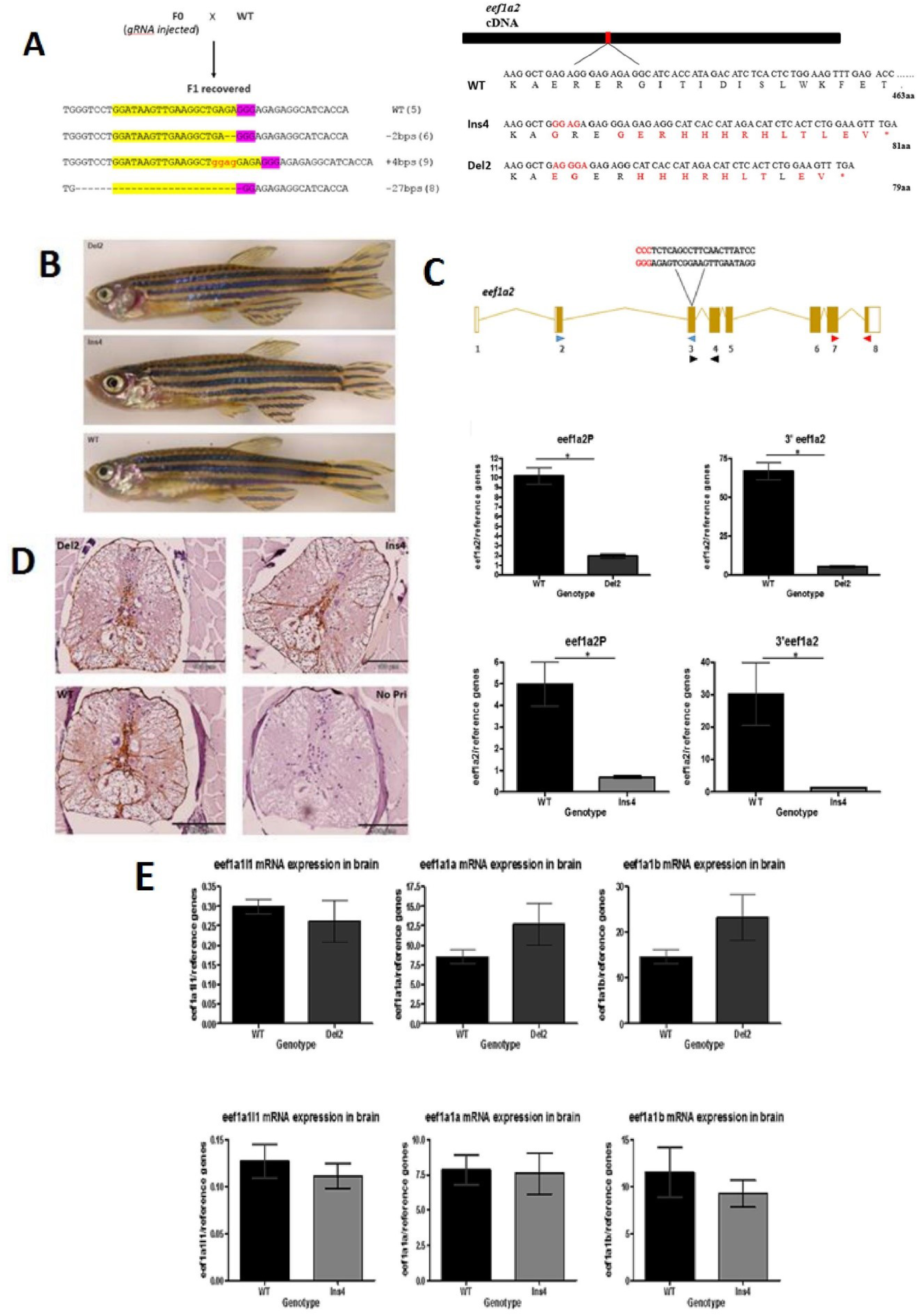
**a** Schematic showing outcross mating of founder (F0) fish and wild-type fish showing recovered F1 sequences (number of F1 fish for each allele are indicated in brackets, target sequences (yellow highlight) and PAM site (purple) with red showing inserted bases) and the predicted effect of Ins4 and Del2 mutant allele with aberrant residues shown in red (right). **b** No overt difference in homozygous Del2 (6 months) and homozygous Ins4 (8 months) adult fish from wild-type (6 months) adult fish. **c** The position of the three different primer sets; eef1a2S (Blue triangle), eef1a2P (black triangle) and 3’eef1a2 (red triangle) is illustrated in relation to the gRNA target site (PAM site sequence in red). Expression levels of *eef1a2* in homozygote Del2 (top panel) and homozygote Ins4 (bottom panel) fish using eef1a2P and 3’eef1a2 primer sets. Results were normalised to *ATPsynth, NADH* and *16S*. (means ± S.E.M; N=3) *p < 0.05. **d** Anti-GFAP antibody stained transverse sections of spinal cords of homozygous Del2 and Ins4 adult fish showed no sign of neurodegeneration. Negative control of a no primary (No Pri) was included which showed no staining. Scale bar = 100µm **e** expression levels of *eef1a1l1* (left), *eef1a1a* (middle) and *eef1a1b* (right) mRNA in homozygous Del2 (top panel) and Ins4 (bottom panel) adult brain. Date are normalised to *ATPsynth, NADH* and *16S* and were presented as means ± S.E.M.; n=3.

#### Expression analysis shows reduced eef1a2 transcripts in Ins4 and Del2 lines

We then went on to investigate the effect of the mutant alleles in each of the Ins4 and Del2 lines. The expression level of *eef1a2* mRNA was assessed using two different set of primers, eef1a2P and 3’eefla2, for both Ins4 and Del2 lines and a third primer set, eef1a2S, for Ins4 only (supplementary figure 2). The position of the primers in relation to the target site is shown in figure 4c.The primer 3’eefla2 was designed such that it is located towards the extreme 3’ end of the *eef1a2* mRNA such that any transcripts downstream of the target site that could lead to translation of a protein would be detected. A marked decrease of *eef1a2* mRNA was seen in both Ins4 and Del2 lines compared to their wild type siblings using each of the different sets of primers (Fig. 4c). Approximately 81% and 92% reduction in the levels of *eef1a2* expression was seen in Del2 homozygous adult brains when compared to their wild-type siblings and a reduction of approximately 86% and 95% in brain tissues of homozygous Ins4 adult fish was seen compared to wild-type. These results suggest that each of the mutant alleles, Ins4 and Del2, lead to decreased messenger RNA levels possibly through nonsense-mediated decay (NMD). Homozygous mutant fish for both lines are thus effectively *eef1a2*-null.

#### Immunohistochemical assessment of Ins4 and Del2 mutants

Complete loss of eEF1A2 in mice has been well characterised and causes motor neuron degeneration of the anterior horn of the spinal cord and muscle wasting, which is of neurogenic origin (Doig *et al.*, 2013). We examined spinal cord sections from homozygous Ins4 and Del2 adult fish to evaluate them for the presence of gliosis (a reactive response to injuries such as neurodegeneration in the central nervous system) using immunohistochemistry for glial fibrillary acidic protein (GFAP). High levels of GFAP staining is seen in human cases of motor neuron degeneration and in animal models, including in the anterior horn of the spinal cord of eEF1A2-null mice (Newbery *et al.*, 2005). However, no increased staining of GFAP was seen in the spinal cord sections of homozygous Del2 and Ins4 mutants when compared to wild-type (Fig. 4d). This result, together with apparently normal histology and normal survival, demonstrates that there is no evidence of neurodegeneration in the spinal cord of either *eef1a2*-mutant zebrafish lines.

#### Other eef1as mRNA remained unchanged in Ins4 and Del2 lines

We next quantified the mRNA level of the other zebrafish *eef1a* genes in order to address both whether the lack of phenotype was a result of a compensatory mechanism from the other genes and also whether any off-target effect involving homologous *eef1a* genes had occurred. The results obtained showed no significant change in the mRNA level of any of the other *eef1a* genes in the brain of adult homozygous Ins4 and Del2 fish (Fig. 4e), suggesting that they were unaffected by the CRISPR/Cas9 targeting of *eef1a2*, and also that no compensatory mechanism had occurred, at least at the mRNA level.

## Discussion

Four different eEF1A genes, namely *eef1a1l1, eef1a1a, eef1a1b* and *eef1a2* are present in the zebrafish genome and share high sequence identity both at the nucleotide (75 - 83 %) and amino acid (89 – 97 %) levels. The nucleotide and amino acid sequences of *eef1a1a* and *eef1a1b* are more similar to each other than the other paralogous genes, suggesting that they arose from the additional teleost-specific genome duplication event which took place at the base of the teleost fish evolutionary lineage (Christoffels et al., 2004). It is clear from our phylogenetic and sequence alignment analyses that the zebrafish Eef1a1a and Eef1a1b are co-orthologues of mammalian eEF1A1. Whilst Eef1a1l1 appears to lack a mammalian orthologue, the data also give strong support to zebrafish Eef1a2 being the sole orthologue of mammalian genes encoding eEF1A2.

We have also demonstrated that all four zebrafish *eef1a* genes are actively transcribed and are expressed in a developmental-specific pattern, consistent with the pattern seen in other vertebrates. During embryogenesis, *eef1a1l1* is the only gene shown to have maternal contribution in addition to zygotic expression as it was detected at all embryonic stages analysed. Interestingly, *eef1a1l1* (formerly referred to as *eef1a*) has been shown to be an essential gene required for early embryonic development in zebrafish (Amsterdam et al, 2004). In this study, mutation in *eef1a1l1* was shown to result in abnormal phenotypes such as small head and eyes from 2 dpf and eventually death at 5 dpf from failure of the swim bladder to inflate. We found that the next *eef1a* genes to be detected were *eef1a1a* and *eef1a1b*, while *eef1a2* was the last to be expressed, at 48 hpf. The detection of *eef1a2* at a later developmental stage is consistent with that of mammals, where its expression is observed much later in development than eEF1A1, gradually replacing it in skeletal muscle and neurons (Knudsen *et al.*, 1993; Lee, Wolfraim and Wang, 1993; Chambers, Peters and Abbott, 1998; Svobodová e*t al.*, 2015).

In all adult tissues analysed, we detected mRNA derived from the *eef1a1l1, eef1a1a* and *eef1a1b* genes. On the other hand, *eef1a2* showed a tissue-specific expression pattern as its mRNA was not present in the liver and was only just detected in the intestine, again similar to the expression seen in mammals. The difference between zebrafish *eef1a2* and the mammalian and *Xenopus* eEF1A2 orthologues however, is the presence of zebrafish *eef1a2* mRNA in spleen and ovary tissue samples. Whilst the expression pattern of *eef1a1a* and *eef1a1b* is in contrast to that of their mammalian orthologues, it is consistent with that of *Xenopus* where eEF1A1 mRNA, in addition to eEF1A2, was detected in adult muscle (Newbery *et al*, 2011). Despite the *eef1a* genes being co-expressed in the tissues, quantification of their expression levels in the brain, muscle and liver suggests that they are not present in equal amounts. As a whole, *eef1a1l1* transcripts are the most abundant, in muscle and liver compared to the other *eef1a* genes, while the levels of all the *eef1a* mRNA species were the same in the brain. The *eef1a2* transcript was the second most abundant in the muscle. In line with being co-paralogues, *eef1a1a* and *eef1a1b* exhibited the same expression pattern in these tissues. The finding that the zebrafish *eef1a* genes display distinct expression profiles suggest they may have evolved unique roles hence their being retained after the duplication events.

We were unable to establish specific expression patterns for zebrafish eef1a genes at the protein level as all commercially available eEF1A antibodies we tested failed to distinguish between the products of the different zebrafish eef1a genes. We were, however, able to see strong expression of *eef1a2* specifically in brain.

We went on to generate two *eef1a2* mutant lines, Ins4 and Del2, using CRISPR/Cas9 genome editing. Our qPCR data showed a substantial decrease in *eef1a2* expression in each of the two mutant lines with either of two sets of primers. This finding indicates that the mutant transcripts are likely targets of nonsense-mediated decay, suggesting that both the Ins4 and Del2 mutations are effectively null, since any remaining transcript would not encode a functional protein. Homozygous loss of *eef1a2* was not lethal in either of our zebrafish mutant lines, in contrast to the situation in mice, which die before 4 weeks in the absence of eEF1A2. Adult homozygous Ins4 and Del2 mutants showed no obvious phenotypic abnormalities, were fertile and produced viable embryos. There are three immediate possible explanations for this discrepancy. Firstly the regenerative capacity of the zebrafish CNS could be masking any neurodegeneration. Secondly, it remains possible the Ins4 and Del2 mutants retain some degree of residual function of *eef1a2*. However, the consistency of the *eef1a2* reduction observed with three different set of primers at different locations from the target site makes this explanation unlikely, and any protein produced from the residual transcripts would be so truncated that they would be highly unlikely to be functional. The third explanation, which seems most likely, is that there is functional redundancy as a result of the three additional *eef1a* genes which were found to be co-expressed with *eef1a2* at the mRNA level. During the course of this work, Cao *et al*. 2017 reported that knockdown of *eef1a2* with morpholinos resulted in abnormal phenotypes including small head size, cardiac failure and skeletal muscle weakness in 2 dpf morphants. Since the development of efficient genome editing techniques in zebrafish it has been increasingly recognised that CRISPR-induced mutants can fail to replicate morpholino-induced phenotypes (Kok et al, 2015). Cao *et al*, did not investigate whether other *eef1a* gene(s) had been down-regulated by the morpholinos used, and it is thus likely that the phenotypes observed were not specific to *eef1a2* but rather the combined effect of the knockdown of one or more of the other *eef1a* genes. Interestingly, a small head, which was one of the phenotypes observed in the *eef1a2* morphants, was also reported in 2 dpf *eef1a1l1* mutant recovered from a large retroviral-mediated insertional mutagenesis screen (Amsterdam *et al*. 2004). Although the phenotypes observed were consistent between different types of morpholinos, translational and splice-site targeting, it is still possible that they may have been the result of a common off-target toxic effect induced by both morpholinos. This type of situation has been demonstrated in the study by Robu *et al.*, 2007, who observed that these two different types of morpholinos induced off-target effects mediated through p53 activation in the zebrafish embryo. Furthermore, Kok et al showed that many morpholino-induced phenotypes in zebrafish, even those that could be rescued by co-injecting with the wild-type mRNA, were likely due to off-target effects and that off-target phenotypes induced by the use of morpholinos occurred much more frequently than was previously thought (Kok et al 2015). The discrepancy between our findings and those of Cao *et al*, 2017 might be due to the different approaches used which in turn induced different responses to *eef1a2* inactivation in zebrafish. Rossi *et al.*, 2015 demonstrated that genetically induced severe mutations resulted in compensatory upregulation of specific proteins which rescued the phenotypes observed. However, our qPCR results show that no upregulation of any of the other *eef1a* genes occur at the mRNA level. Again, this is consistent with the idea of functional redundancy of *eef1a* genes as the most likely explanation for the lack of abnormalities after the loss of *eef1a2* in Ins4 and Del2 mutant lines. In contrast to morpholinos, CRISPR/Cas9 has been shown to have negligible off-target effects in zebrafish. Using next-generation sequencing (NGS), Hruscha *et al.*, 2013 demonstrated that off-target effects were limited in founder fish. This study was small but was supported by another larger study in which the target sites for five gRNAs targeted to different genes were analysed. One 3 base pair deletion was found, in only one of the 25 off-target loci tested (Varshney *et al.*, 2015). The possibility of random off-target events occurring in the Ins4 and Del2 lines cannot be ruled out, but, the use of mutant fish starting from the F2 generation and resulting from an outcross of the F0 fish with wild-type fish should minimise the risks as off-target mutations should segregate away from the Del2 and Ins4 *eef1a2* mutation (Schulte-Merker and Stainier, 2014). Furthermore, the comparison of two independent mutations and the lack of any observable abnormalities suggest that off-target effects are not a concern.

Overall, our results suggest that ablating the expression of eEF1A2 in zebrafish is unlikely to provide a model system in which to study disease-causing loss of function mutations. However, if the epilepsy-causing missense mutations seen in humans in fact represent a toxic gain of function, our new Del2 and Ins4 lines could provide an important resource in which to test the effects of expression of mutant eEF1A2 in the form of human mRNA.

## Acknowledgements

We are grateful to staff of the MRC HGU zebrafish facility for their technical support, Witold Rybski for assistance with the microinjection and zebrafish dissection training, Zhiqiang Zeng for his kind gift of Cas9 mRNA and Liz Patton group for their kind support and helpful advice.

## Supplementary figures

**Supplementary figure 1.**
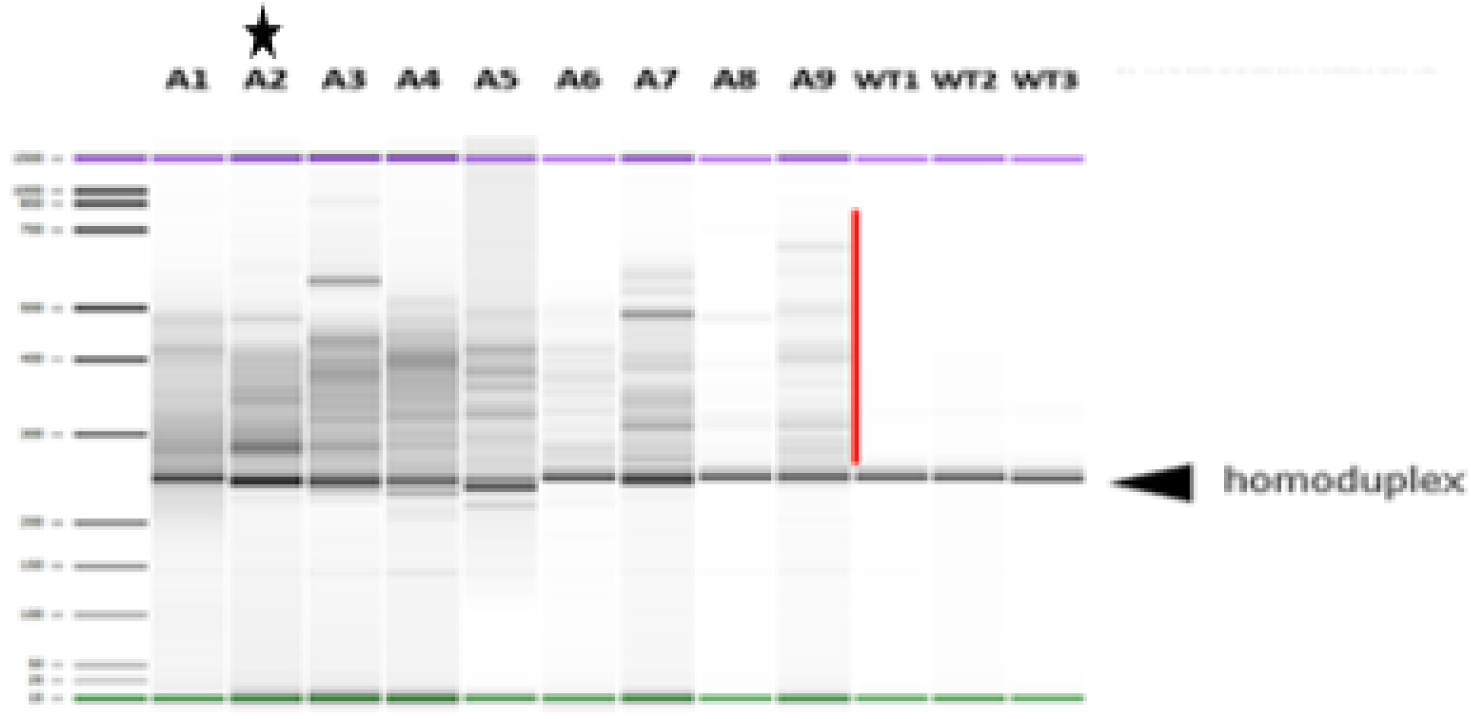
Screening of potential F0 mutants injected with gRNA by running PCR amplicons of target site on the Agilent 2100 Bioanalyser. A mismatch between wild type and mutant strands gives rise to heteroduplex ((shown by the red line)), which indicates the presence of indels in these fish which are mosaic at this stage. Black star indicates founder (F0) fish used to generate Ins4 and Del2 lines. WT1, WT2 and WT3 indicate PCR products obtained from three different uninjected wild-type fish fin-clippings.

**Supplementary figure 2.**
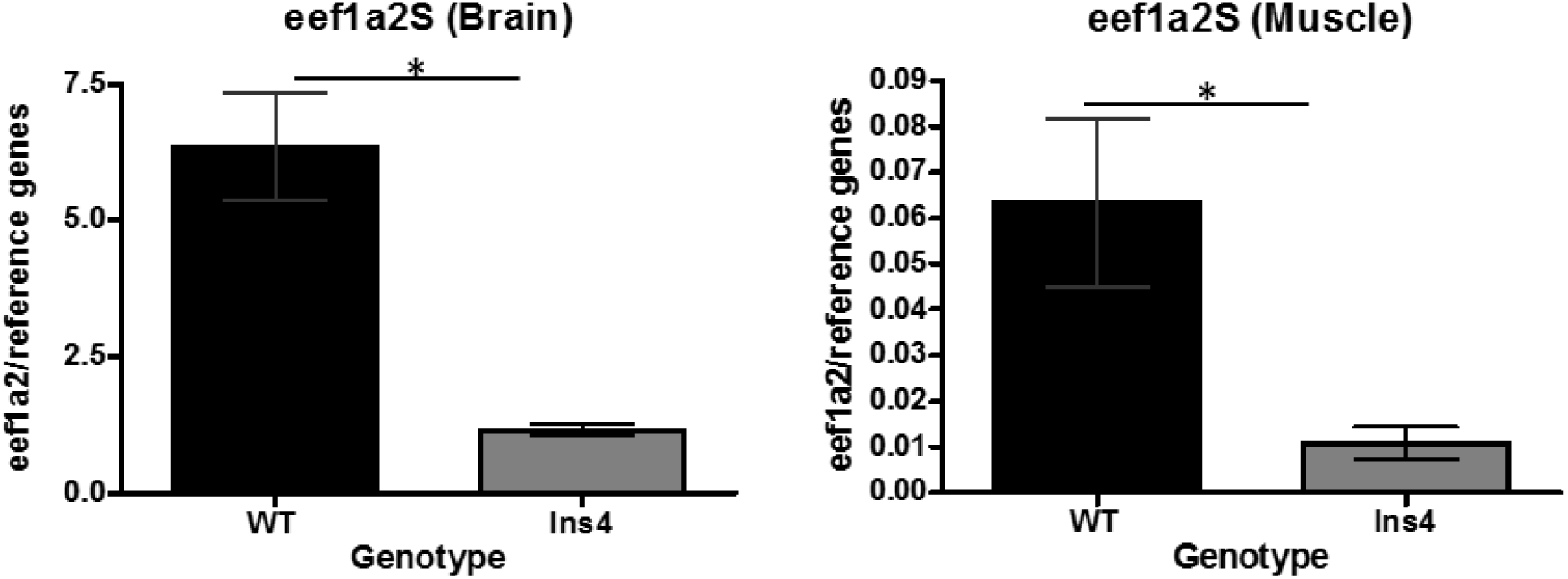
Analysis of *eef1a2* transcripts in Ins4 mutants using eef1a2S. Reduced *eef1a2* transcript levels in F2 Ins4 homozygous (3 months) brain and muscle tissues was also noted using this set of primers.

